# Spike desensitisation as a mechanism for high-contrast selectivity in retinal ganglion cells

**DOI:** 10.1101/2022.08.11.503581

**Authors:** Le Chang, Yanli Ran, Olivia Auferkorte, Elisabeth Butz, Laura Hüser, Silke Haverkamp, Thomas Euler, Timm Schubert

**Author notes:** Corresponding authors: Thomas Euler, Timm Schubert. Submitting author: Timm Schubert. Competing interests: The authors declare no competing financial interests.

## Abstract

In the vertebrate retina, several dozens of parallel channels relay information about the visual world to the brain. These channels are represented by the different types of retinal ganglion cells (RGCs), whose responses are rendered selective for distinct sets of visual features by various mechanisms. These mechanisms can be roughly grouped into synaptic interactions and cell-intrinsic mechanisms, with the latter including dendritic morphology as well as ion channel complement and distribution. Here, we investigate how strongly ion channel complement can shape RGC output by comparing two mouse RGC types, the well-described ON alpha cell and a little-studied ON cell that is EGFP-labelled in the *Igfbp5* mouse line and displays an unusual selectivity for high-contrast stimuli. Using patch-clamp recordings and computational modelling we show that in ON *Igfbp5* cells – but not in the ON alpha cells – a higher activation threshold and a pronounced slow inactivation of the voltage-gated Na^+^ channels are responsible for the distinct contrast tuning and transient responses of ON *Igfbp5* RGCs, respectively. This study provides an example for the powerful role that the last stage of retinal processing can play in shaping RGC responses.

**SIGNIFICANCE STATEMENT:** Here, we investigated, how voltage-gated sodium channels contribute to shaping the light responses of mouse retinal ganglion cells. Using single-cell electrophysiology and computational modelling, we studied a ganglion cell type that displays highly transient responses and an unusual selectivity for visual high-contrast stimuli. We found that the cell’s characteristic responses were largely determined by intrinsic mechanisms, notably, a high activation threshold and a pronounced slow inactivation of its voltage-gated sodium channels. Therefore, our study demonstrates how sodium channels at the last stage of retinal signal processing can contribute to shape retinal output to higher visual areas the brain; it also adds a rare example for how channel complement can be directly linked to feature selectivity.

## INTRODUCTION

Retinal ganglion cells (RGCs) and their upstream circuits detect and encode specific visual features and relay this information along parallel pathways to higher visual centres in the brain (reviewed in (Kerschensteiner, 2022)). Functional, anatomical, and genetic evidence (Baden et al., 2016; Bae et al., 2018; Goetz et al., 2021; Rheaume et al., 2018) support the presence of at least 40 RGC types in the mouse retina.

RGCs receive their excitatory drive mostly from bipolar cells (BCs), which relay the photoreceptor signal to the inner retina, and their inhibitory input from amacrine cells (ACs) (reviewed in (Diamond, 2017)). As the dendrites of different RGC types arborize at distinct IPL depths (Bae et al., 2018; Helmstaedter et al., 2013; Sümbül et al., 2014), they pick up inputs from distinct sets of BC and AC types (Field et al., 2010; Helmstaedter et al., 2013). The selective connectivity with presynaptic neurons in the inner plexiform layer (IPL) is considered the foundation of the feature-selectivity of RGC pathways (e.g. (Helmstaedter et al., 2013; Neumann et al., 2016; Roska and Werblin, 2001; Yu et al., 2018)).

The response properties of an RGC type are typically determined by a hierarchy of mechanisms. For instance, the temporal response of an RGC to a light-step is shaped by presynaptic circuit components, such as glutamate receptor kinetics along the BC-RGC pathway (Awatramani and Slaughter, 2000; DeVries, 2000; Turner and Rieke, 2016; Yu et al., 2018), as well as by AC input (Asari and Meister, 2012; Cui et al., 2016; Franke et al., 2017; Jacoby et al., 2015; Nikolaev et al., 2013; Nirenberg and Meister, 1997). Moreover, the spatial RF is jointly formed by horizontal cells (Drinnenberg et al., 2018) and ACs (Diamond, 2017) in the outer and inner retina, respectively. In particular, AC circuits are very versatile “function modifiers”: For example, in the highly contrast-sensitive On alpha RGCs (Krieger et al., 2017), different AC circuits converge to provide On and Off inhibition to balance tonic excitatory drive from BCs (Park et al., 2018; Sawant et al., 2021); in On delayed RGCs, they provide a fast, excitatory surround through disinhibition (Mani and Schwartz, 2017).

In addition, RGC types differ in their expression of ion channels (Rheaume et al., 2018; Siegert et al., 2012) and how they are distributed across the cell (reviewed in (Van Hook et al., 2019)). This ion channel complement, in combination with the RGC’s specific dendritic geometry, determines how synaptic input is integrated (Poleg-Polsky and Diamond, 2011; Ran et al., 2020; Schachter et al., 2010) and how the resulting signal is translated by the cell’s spike generator into action potentials on the optic nerve (Kim and Rieke, 2001, 2003; Mobbs et al., 1992; Raghuram et al., 2019; Werginz et al., 2020; Wienbar and Schwartz, 2022). Hence, RGCs “themselves” can importantly transform the input they receive from their circuits and thereby shape the retina’s output to the brain (reviewed in (Branco and Häusser, 2010; Stuart and Spruston, 2015; Tran-Van-Minh et al., 2015)).

The contribution of the aforementioned mechanisms to the RGC output varies among RGC types, yielding the diversity of feature representations, such as local edges (Levick, 1967; van Wyk et al., 2006; Zhang et al., 2012), approaching objects (Münch et al., 2009), “uniformity” (Jacoby et al., 2015; Levick, 1967; Tien et al., 2016), and direction of motion (Barlow et al., 1964). For this, intrinsic properties may play a pivotal role: For instance, On-Off direction-selective (DS) RGCs employ a range of mechanisms, including directionally-tuned inhibitory and likely excitatory input from asymmetrically wired ACs, but their dendritic morphology and active channel distribution substantially contribute to these cells’ DS tuning (reviewed in (Borst and Euler, 2011; Mauss et al., 2017)).

Here, we study the role of a mechanism at the very end of the signal-shaping hierarchy, the encoding of the RGC membrane voltage into spike trains. We performed electrical single-cell recordings from a transient On RGC type that is EGFP-labelled in the transgenic *Igfbp5* mouse line (Siegert et al., 2009), which likely corresponds to G_18a_ (“ON trans”; (Baden et al., 2016)), “ON transient small RF” (Goetz et al., 2021), and “6sn” (Bae et al., 2018). These cells display an unusual non-linear response behaviour in that they are sharply tuned to high-contrast light stimuli. Using single-cell current analysis and computational modelling, we show that desensitisation of the *Igfbp5-*positive transient On small (tOn-small) RGC*’*s intrinsic spike generator accounts for both the transience of their light response and their selectivity for high contrasts.

## RESULTS

### EGFP-expressing neurons in the Igfbp5 mouse retina

Transgenic mice, in which specific RGC types are fluorescently labelled, greatly facilitate investigating RGC function (e.g. (Bleckert et al., 2014; Münch et al., 2009; Rousso et al., 2016; Yao et al., 2018)). While screening the Gene Expression Nervous System Atlas (GENSAT) database of transgenic mice for selective lines (Siegert et al., 2009), we were struck by the very distinctive pattern of retinal EGFP expression in the *Igfbp5* (insulin-like growth factor-binding protein 5) mouse line (Fig. 1A,B). In the *Igfbp5* retina, the EGFP is expressed in two prominent, equally thick bands of processes along IPL sublaminae 2 and 3 (Fig. 1A, Fig 1-1A-C), extending just in between the two choline acetyltransferase (ChAT) -positive bands (Fig. 1-1D,E). This pattern is reminiscent of the glypho (glycogen phosphorylase) staining in the macaque monkey retina (Majumdar et al., 2008). We detected EGFP in a subset of BCs and ACs in the inner nuclear layer (INL; Fig. 1A; for details, see Fig. 1-1A-E), as well as in some RGCs and displaced ACs in the ganglion cell layer (GCL; Fig. 1B; Fig. 1-1F).

**Figure 1.**
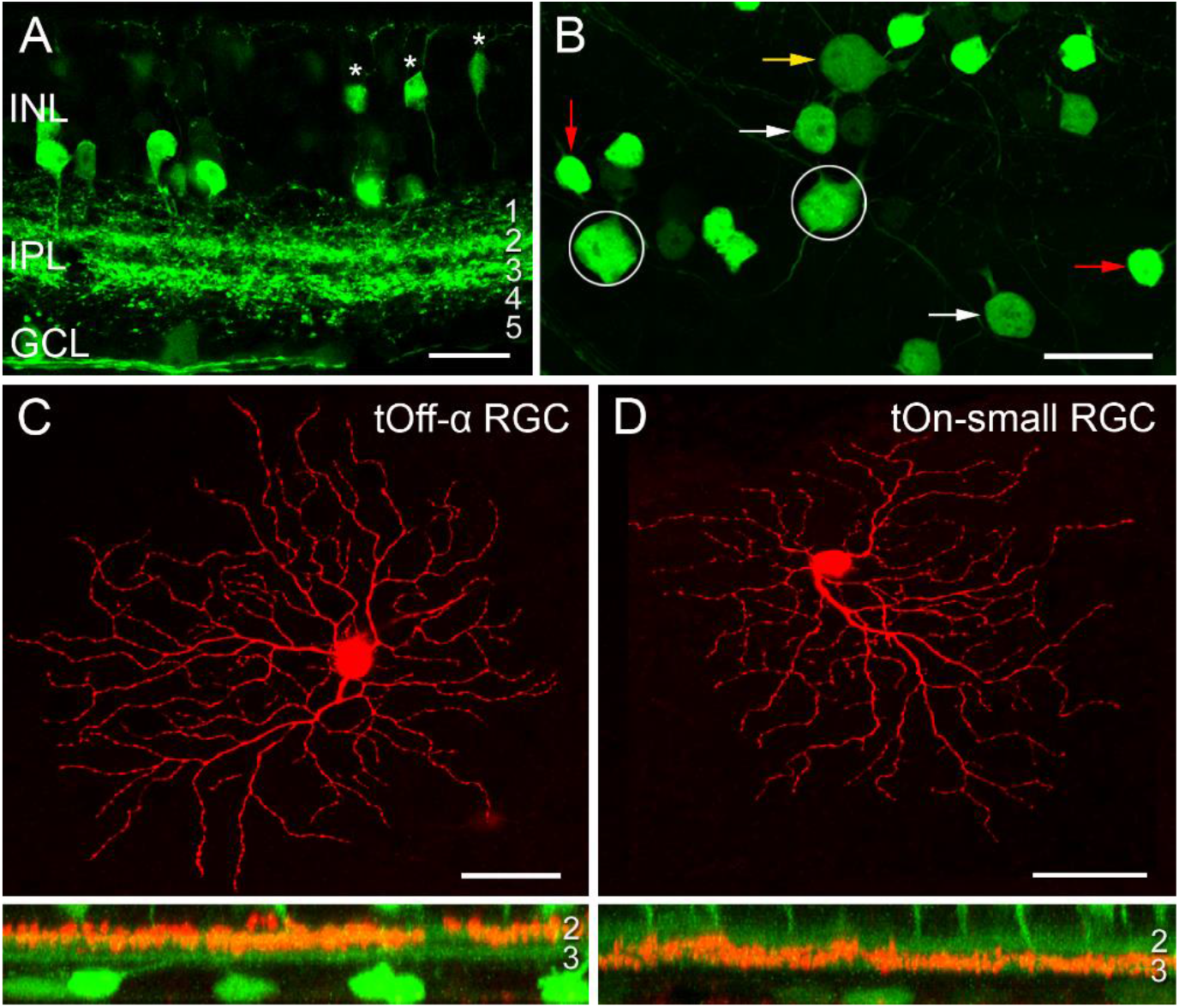
EGFP-labelled cells in the Igfbp5 transgenic mouse line. **A**, EGFP expression in a vertical section of the Igfbp5 retina, with EGFP expressed in bipolar cells (BCs, asterisks) and amacrine cells (ACs) in the inner nuclear layer (INL), as well as in displaced amacrine cells and retinal ganglion cells (RGCs) in the ganglion cell layer (GCL). EGFP-labelled processes extend along sublaminae 2 and 3 of the inner plexiform layer (IPL). **B**, Whole-mount showing mosaic of EGFP-expressing somata in the GCL. Examples of putative displaced ACs (red arrows), Off RGCs with medium-sized (white arrows) and large (yellow arrow) somata, and On RGCs (circles) indicated. **C**,**D**, Examples of EGFP-positive RGCs: tOff-α RGC (C), and tOn-small RGC (D) filled with Neurobiotin during electrical recording. Below each panel, z-projections of the cell’s dendritic arborisation. Scale bars: A, 20 µm; B, 25 µm; C and D, 50 µm.

In the GCL, four cell types could be distinguished based on their soma size and labelling intensity (Fig. 1B): very bright cells with small somata (soma diameter: 9.1 ± 0.5 µm, mean ± s.d., n = 29), dimmer cells with medium-sized (12.0 ± 0.5 µm, n = 24) or larger somata (14.8 ± 0.9 µm, n = 34), and very dim cells with large somata (17.5 ± 0.7 µm, n = 14).

Dye-injections (n = 40) revealed that the small, brightly labelled cells were likely displaced wide-field ACs (Lin and Masland, 2006): They featured 6-10 primary dendrites that sometimes bifurcated close to the soma (∼14 dendrites/cell in total), rarely crossed each other and narrowly stratified in IPL sublamina 3 (Fig. 1-1F). The dendritic fields of neighbouring cells overlapped frequently, resulting in dense retinal coverage (Fig. 1-2). These putative ACs closely resembled glypho-positive On ACs in the macaque (Majumdar et al., 2008).

The three other *lgfbp5-*positive cells in the GCL were monostratified RGCs (Fig. 1C,D; Fig. 2-1A-C_) with their dendrites in either one of the two EGFP bands (*cf*. Table 1). The cells with the largest, dimmest somata were transient Off-alpha (tOff-α) RGCs (Fig. 1B,C; Fig. 2-1B) (Krieger et al., 2017; Ran et al., 2020; van Wyk et al., 2009), as confirmed by immunolabeling (Fig. 2-1E-H) for markers such as SMI32 (Fig. 2-1E,F), which strongly labels sustained On alpha (sOn-α) and tOff-α cells in mouse (Bleckert et al., 2014; Coombs et al., 2006). The remaining two labelled cell populations had dendritic arbour diameters of around 200 µm.

**Table 1.**
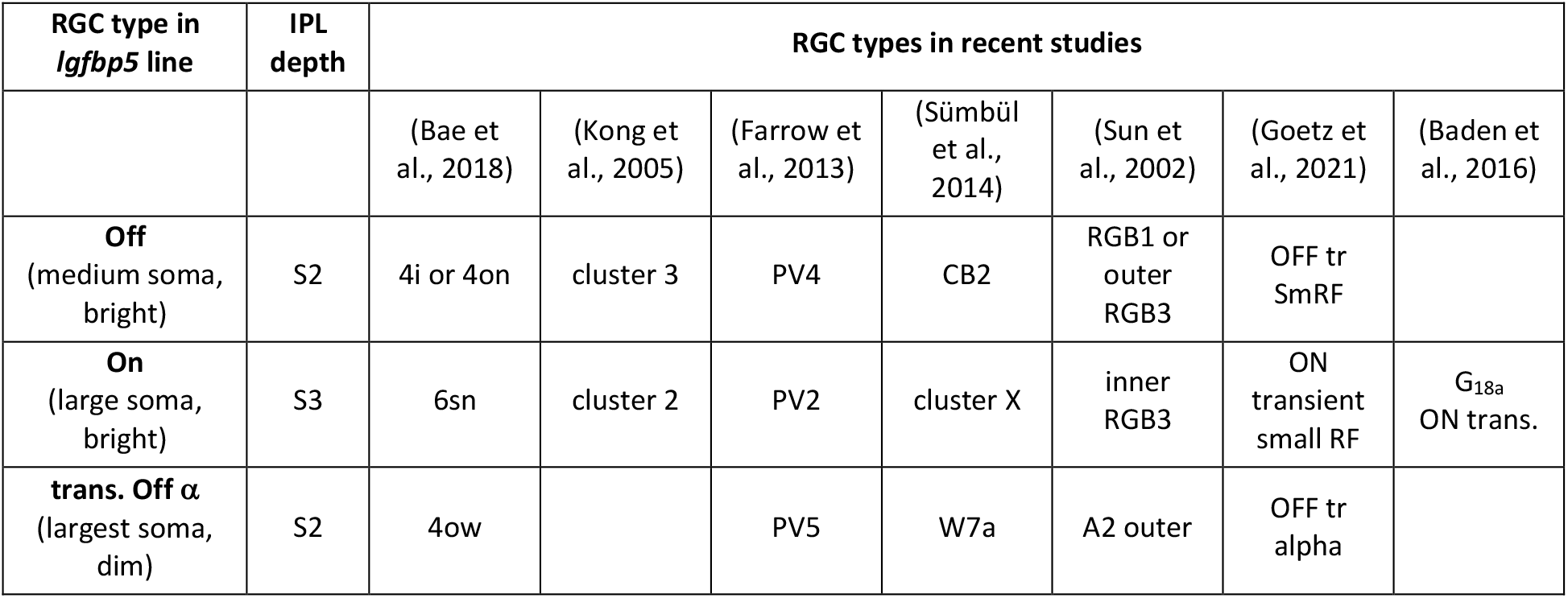
Igfbp5-positive RGCs and their presumed morphological counterparts in earlier mouse studies. Missing entries indicate that type assignment is unclear.

The *lgfbp5-*positive cells with medium-sized somata were presumably Off cells, as their dendrites stratified in sublamina 2; their morphology (Fig. 2-1A) resembled that of “4i” or “4on” cells described in (Bae et al., 2018), cluster 3 (Kong et al., 2005), PV4 (Farrow et al., 2013), CB2 (Sümbül et al., 2014), and RGB1 or “outer” RGB3 cells (Sun et al., 2002).

The mouse *lgfbp5-*positive RGCs with the larger somata (Fig. 1D and 2A,C; Fig. 2-1C) were presumably On cells as they stratified in sublamina 3; their morphology resembled that of “6sn” (Bae et al., 2018), “ON transient small RF” (Goetz et al., 2021), cluster 2 (Kong et al., 2005), PV2 (Farrow et al., 2013), „cluster X” (Sümbül et al., 2014), and “inner” RGB3 cells (Sun et al., 2002). For simplicity, we refer to the *lgfbp5*-positive On RGCs in the following as transient On small (tOn-small) cells.

**Figure 2.**
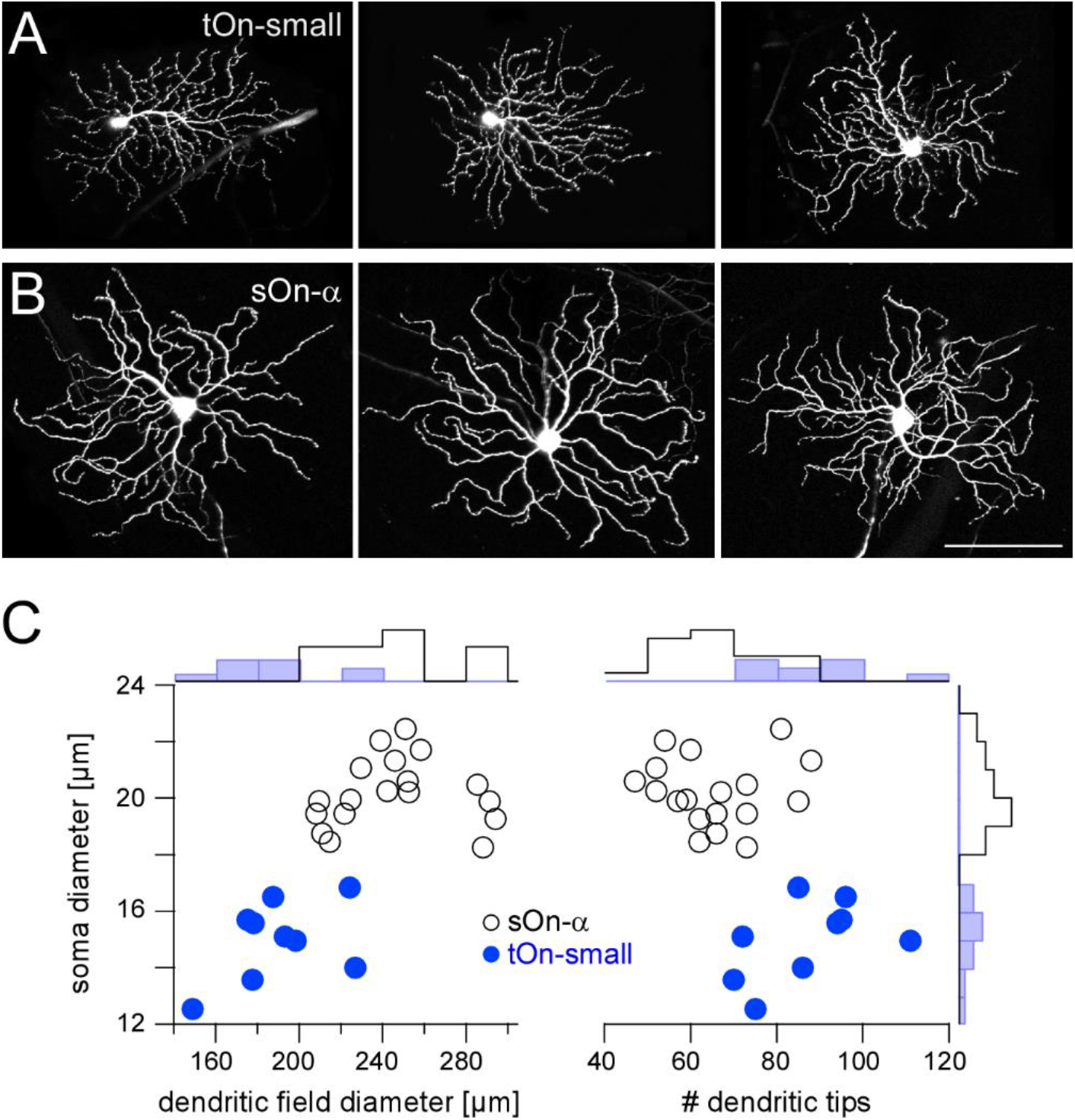
Igfbp5-positive tOn-small ganglion cells differ morphologically from sustained sOn-α cells. **A**,**B**, Examples of dye-injected RGCs: three tOn-small (A) and three sOn-α cells (B). To avoid confounds related to morphological variability across the retina (Bleckert et al., 2014), the cells were collected (and recorded) from the ventral retina (∼0.8 mm from the optic disc). **C**, Soma diameter vs. dendritic field diameter (left) and vs. number of dendritic tips (right) for the two RGC types, with distribution histograms at the sides. Scale bar: A,B, 100 µm.

Mouse transient On-alpha (tOn-α) RGCs stratify at the same level as the tOn-small cells, but have a slightly larger dendritic tree (Goetz et al., 2021) and are osteopontin-positive (Krieger et al., 2017). We think that the tON-small RGCs are homologous to primate parasol cells (Fig. 2-1I) – as was suggested for cluster 12 cells of Farrow and Masland (2011).

### tOn-small RGC responses are highly transient

When we characterized light-evoked signals in tOn-small cells using two-photon-guided electrical recordings (Methods), we were intrigued by their very transient responses – with the spike rate increasing almost instantaneously at light-onset but then quickly dropping to zero within ∼200 ms and no response at light-offset (Fig. 3-1A. As such transient responses were described in a type of direction-selective On RGC in rabbit (Kanjhan and Sivyer, 2010), we recorded tOn-small cell responses to a moving bar stimulus, but did not find any substantial directional tuning in these cells (Fig. 3-1B; DSi = 0.043 ± 0.019, n=5 cells).

In the following, we studied the mechanisms underlying the characteristic transient responses of tOn-small cells in the mouse retina. For comparison, we recorded the well-described sOn-α RGCs (Peichl et al., 1987), which are known for their sustained responses and high contrast sensitivity (Bleckert et al., 2014; Krieger et al., 2017; Pang et al., 2003; Schwartz et al., 2012; van Wyk et al., 2009) (Fig. 2B). Morphologically, the two RGC types could be distinguished easily, with tOn-small cells having smaller somata and dendritic fields, but more dendritic tips than sOn-α cells (Fig. 2).

### tOn-small cells encode high-contrast signals

First, we studied the spatial receptive field (RF) organisation of tOn-small cells using spot stimuli with varying diameters centred on the soma and sinusoidally modulated at 1 Hz (Fig. 3A,B), in comparison to sOn-α cells. Using spectral analysis, we then calculated the amplitude of the cells’ fundamental response component (Methods). With increasing spot diameter, amplitudes first increased and then declined (Fig. 3B), indicative of centre-surround antagonism. To quantify the spatial RFs, we used a Difference-of-Gaussians (DOG) model to interpret the recorded area summation data (Methods). Consistent with their larger dendritic field diameters (Fig. 2), sOn-α cells had larger RF centres than tOn-small cells (Fig. 2C).

**Figure 3.**
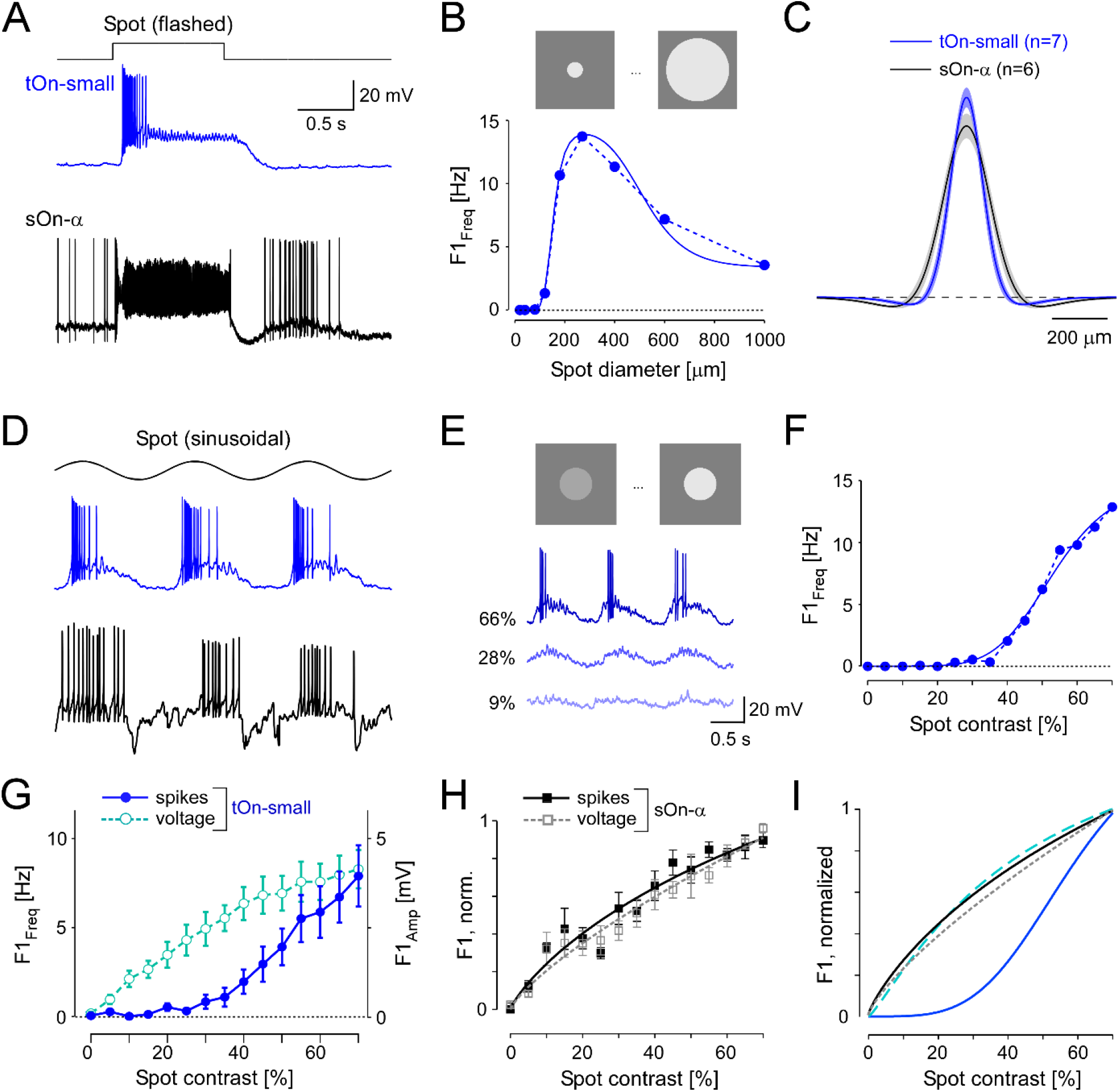
tOn-small ganglion cells have smaller receptive fields (RFs) and higher contrast thresholds compared to sOn-α cells. **A**, Intracellularly recorded voltage responses of an tOn-small (blue) and a sOn-α (black) to a 1-s flashed spot (200 µm in diameter). **B**, Fundamental (F1) spiking response (F1_Freq_ in Hz) as a function of spot diameter (1 Hz, 57% mean contrast) for the same tOn-small cell as in (A). **C**, Mean and SEM of estimated RF profiles, using difference-of-Gaussians (DOG) RF model for n=7 tOn-small and n=6 sOn-α cells (Methods). **D**, Responses of the same cells as in (A) to a 1-Hz sinusoidal stimulus (270 µm in diameter). **E**, Exemplary responses of an tOn-small cell for different contrasts. **F**, F1_Freq_ as a function of stimulus contrast (D_1_; 270 µm diameter spot; data fit with Naka-Rushton equation; Methods) for the same tOn-small cell as in (E). **G**, Response-contrast functions for tOn-small cells (blue, spikes, n=7 cells; cyan, voltage, n=3 cells). **H**, Normalized response-contrast curves for an additional set of sOn-α cells (black, spikes, n=6 cells; grey, voltage, n=6 cells; Naka-Rushton fits). **I**, Normalized response-contrast curves (dataset from (G,H); Naka-Rushton fits). Symbols and error bars represent mean and SEM, respectively.

In the DOG model, we included a static-nonlinearity to convert the weighted stimulus into a firing rate. Because the area summation data by itself was not sufficient to characterize this nonlinearity, we also presented spots with fixed diameter but with different contrasts (Fig. 3D). Here, we used the spot diameter (270 µm) that elicited the maximal spiking response. Other than sOn-α cells, tOn-small cells were insensitive to weak stimuli: Sinusoidally modulated spots with contrasts lower than 40% hardly evoked any spiking response (Fig. 3E,F).

We also analysed the graded voltage responses recorded in the whole-cell current-clamp mode (with spikes digitally removed, *Methods*) and found that the strong threshold-like behaviour was not reflected in the graded response: tOn-small cells showed a detectable depolarization even to the smallest contrast tested (5%, Fig. 3G, cyan circles vs. blue circles), resulting in a contrast sensitivity curve similar to that for the sOn-α cells’ spiking response (Fig. 3H; black squares). Note that sOn-α cells displayed a very similar contrast sensitivity curve for spikes and graded voltage (Fig. 3H; filled black vs. open grey squares). Together, this suggests that the conversion of voltage into a spiking response differs between tOn-small and sOn-α cells (Fig. 3I). A possible explanation may be a larger difference between resting potential and spiking threshold in tOn-small vs. sOn-α cells. While the resting potential (V_rest_) of tOn-small cells (−66.8 ± 3.4 mV; n=6) indeed was slightly more hyperpolarized than that of sOn-α cells (−64.8 ± 3.3 mV; n=6), the difference seems too small to explain the observed difference in contrast sensitivity alone.

The mismatch in contrast sensitivity between spiking and graded voltage responses in tOn-small cells is striking. Related to this was our finding that when stimulated with a bright flash, the cells displayed a sustained graded voltage response – in contrast to the transient nature of their spiking response (Fig. 3A, top). We therefore studied temporal processing in tOn-small (and, for comparison, sOn-α) cells with a homogeneous white-noise stimulus (Methods), using again the optimal spot diameter. First, we calculated the spike triggered average (STA) as an estimate of the cell’s temporal linear filter (Chichilnisky, 2001). tOn-small cells had highly biphasic filters, with an On-lobe close to the time of spike and an almost similarly strong Off-lobe further away (Fig. 4A_1_; blue curve). We quantified the filter’s biphasic nature by calculating the ratio between Off- and On-lobe peak (Chander and Chichilnisky, 2001), yielding a biphasicity index (*B*_*i*_) of 0.60 ± 0.19 (n=6) for the spike response of tOn-small cells. In contrast, the linear filter estimated from their graded voltage response (*B*_*i*_ = 0.28 ± 0.17, n=2) and the STA-derived linear filters of sOn-α cells (*B*_*i*_ = 0.15 ± 0.09, n=5) were more monophasic (Fig. 4A_1_; dashed cyan and black curve, respectively).

**Figure 4.**
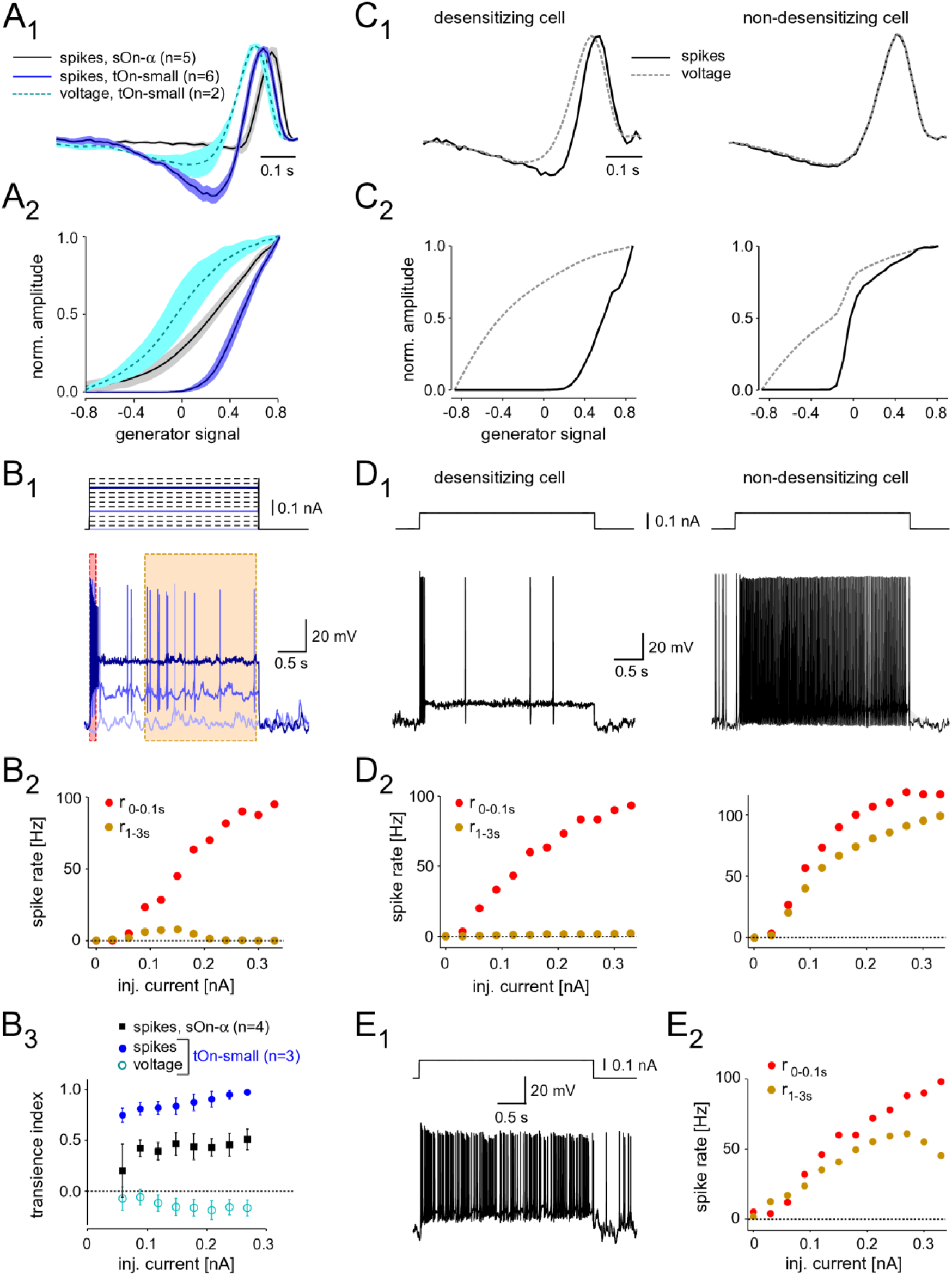
Spike desensitisation in tOn-small ganglion cells accounts for their biphasic temporal filter and high contrast threshold. **A**, Temporal linear filter (A_1_) and static nonlinearity (A_2_) estimated using a white noise stimulus (Methods) for tOn-small (blue, spikes, n=6 cells; cyan, graded voltage responses, n=2 cells) and sOn-α cells (black, spikes, n=5 cells). **B**, Intracellularly recorded voltage responses of a tOn-small cell (B_1_, bottom, resting potential V_rest_ = -66 mV) in response to current injections of varying amplitude (B_1_, top). B_2_, Firing rate of the same cell right after current injection onset (0-0.1 s, red) and at steady-state (1-3 s, brown), as functions of injected current (for time windows, see B_1_). B_3_, Relationship between spike desensitisation and injected current for tOn-small (blue, spikes; cyan, graded voltage responses, n=3) and sOn-α cells (black, spikes, n=4). Spike desensitisation was quantified as transience index (Ti_spike_=1 − (r_1−3_s/r_0−0.1s_)or Ti_V_=1 _− (_V1_−3_s/V_0−0_.1s), with spike rate r and baseline-corrected graded potential V, respectively). Symbols and error bars represent mean and SEM, respectively. **C**, Temporal linear filter (C_1_) and static nonlinearity (C_2_) estimated using simulated responses of two modelled RGCs (left: with strong spike desensitisation; right: with weak spike desensitisation) to a white noise stimulus (for details, see text). **D**, Voltage responses of the two modelled cells to current injection of 0.1 nA (D_1_) and their firing rates as functions of injected current (D_2_, analogous to recording in B_2_). **E**, Voltage recording of a sOn-α cell (V_rest_ = -64 mV) in response to current injections of 0.1 nA (E_1_) and its firing rate as a function of injected current (E_2_, analogous to B_2_).

The difference between linear filters estimated from spike vs. voltage responses indicates that the nonlinearity underlying spike generation in tOn-small cells is not a “trivial” static nonlinearity, i.e. a simple nonlinear function whose output only depends on the present value of input. Because the linear filter estimation is expected to be resistant to a static nonlinearity (Chichilnisky, 2001), the involvement of a static nonlinearity should result in similar filters – independent of the response modality used (spikes vs. graded voltage). To characterize the nonlinearity in tOn-small cells, we calculated the relationship between the linear response (“generator signal” = the convolution of estimated linear filter and stimulus) and the recorded response to the white noise stimuli. The resulting nonlinear function for the spike responses of tOn-small cells had a much higher threshold than that for their voltage responses and the spike responses of sOn-α cells (Fig. 4A_2_).

Many response features of RGCs, including temporal properties and contrasts sensitivity, can be shaped by synaptic interactions (Introduction). However, the clear differences in transiency and contrast sensitivity between spiking and graded voltage responses observed in tOn-small (but not sOn-α) RGCs point at the intrinsic spike generator of tOn-small cells. Therefore, in the following we focused on this final step in retinal signal transformation.

### Spike generation in tOn-small RGCs is readily desensitised by constant current injection

The comparison between spike and graded voltage responses revealed two distinct features of spike generation in tOn-small cells: (*i*) the high-threshold nonlinearity and (*ii*) the pronounced biphasicity of the linear filter. The first feature may simply result if the spike threshold is substantially higher than the resting potential. The second feature implies more complicated nonlinearities, such as spike desensitisation (Goldin, 1999). We tested the spike generator by injecting constant current of different amplitudes into the cells while recording their voltage responses (Fig. 4B_1_). We found that for tOn-small cells, injection of a large current elicited a burst of spikes at injection onset but then spiking stopped rapidly (Fig. 4B_1_, dark blue trace). The initial spike frequency (time window: 0-0.1 s) scaled almost linearly with the amplitude of injected current (Fig. 4B_2_, red symbols), whereas the steady-state spike frequency remained as low as ∼10 spikes/s or less (orange symbols). As a result, transiency of the spike response increased with stimulus strength, as indicated by the increasing ratio between initial spiking response and steady state (Fig. 4B_3_). This increase was absent or less pronounced in graded voltage responses of tOn-small cells and spiking responses of sOn-α cells, respectively. Together, this points at strong desensitisation of the spike generator in tOn-small cells.

### Spike desensitisation model mimics response properties of tOn-small RGCs

We constructed a simple model to explore how varying levels of spike desensitisation affect the signal encoding properties of desensitising and non-desensitizing cells, such as tOn-small and sOn-α cells, respectively (Methods). We modelled responses to the white-noise stimulus by calculating the amplitude of the input current to the cell using a simple linear-nonlinear (LN) model. First, for the desensitising case (Fig. 4C_1,2_; left), we estimated the linear filter and nonlinearity from graded voltage responses measured in tOn-small RGCs (Fig. 4A_1,2_). Using the voltage output of the modelled cell, we then estimated linear filter and nonlinearity for the spike responses. We found that the spike response-derived linear filter was more biphasic than that based on the graded voltage response, very similar to what we measured in tOn-small cells (cf. Fig. 4A_1,2_). For the non-desensitising case (Fig. 4C_1,2_; right), the two linear filters were nearly identical. In both model cells, the nonlinearity for spike responses exhibited a higher threshold than that for graded voltage responses, however, only the threshold in the desensitising cell was higher than the linear response to 0% contrast (generator signal = 0; Fig. 4C_2_; cf. panel A_2_). Therefore, low-contrast stimuli (generator signal ∼0) evoke only sub-threshold responses in the desensitising model cell – very similar to the responses recorded in tOn-small cells (cf. Fig. 3E).

We also used current steps with different amplitudedramatically dropped at 20 Hz already fosr as model input (Fig. 4D). As expected, the desensitising model cell displayed little or no spike response at the steady state (1-3 s) of the current step (Fig. 4D; left), very similar to the responses recorded in tOn-small RGCs (Fig. 4B_1_). In contrast, the steady-state response of the non-desensitising model cell was nearly as strong as its response at the current injection onset (0-0.1s; *r*_2−3 *s*_⁄*r*_0−0.1 *s*_ >70%) for each current amplitude (Fig. 4D, right), very similar to the recorded response of sOn-α cells (Fig. 4E).

In conclusion, our model supports that spike desensitisation and a high spiking threshold may account for two distinct response features of tOn-small RGCs: their strong transience and their selectivity for high contrasts. Note that our simple desensitising model generates responses very similar to those of tOn-small cells, whereas the match between the non-desensitising model and sOn-α cells is not as good: The nonlinearity is “steeper” in the non-desensitising model cell than that estimated from sOn-α cell recordings (Fig. 4C_2_ right vs. 4A_2_). This discrepancy suggests that a sOn-α cell model requires additional properties, such as, for example, a higher noise level in the membrane potential, which can result in a smoother nonlinearity (Anderson et al., 2000).

### Voltage-gated Na^+^ channels in tOn-small RGCs are high-threshold activated

Our modelling predicts that the Na^+^ channel desensitization observed in tOn-small cells may result from differences in voltage-gated Na^+^ channel (VGSC) activation, inactivation and/or recovery from inactivation (Carter and Bean, 2011). To test this experimentally, we recorded Na^+^ currents in these cells and, for comparison, in sOn-α cells (Figs. 5 and 6). To isolate VGSCs, we blocked voltage-gated K^+^ channels with Cs^+^ and TEA in the electrode solution and voltage-gated Ca^2+^ channels with Cd^2+^ in the extracellular solution (Methods). With voltage-step protocols to characterize VGSC activation (Fig. 5A_1_) and inactivation (Fig. 5A_2_) we found VGSCs in tOn-small cells to activate at higher potentials than those in sOn-α cells (Fig. 5A_3_; tOn-small, -51.1 ± 3.9 mV, n=6; sOn-α, -67.5 ± 3.2 mV, n=7; at 5% activation estimated from sigmoidal fit), which explains why a substantial depolarisation (approx. >15 mV from *V*_*Rest*_) is required to trigger spikes in tOn-small cells, consistent with their selectivity for high contrasts. However, the inactivation profiles were almost identical (Fig. 5A_3_), suggesting that at least for this protocol, the VGSCs in the two cell types share similar steady-state inactivation properties.

**Figure 5.**
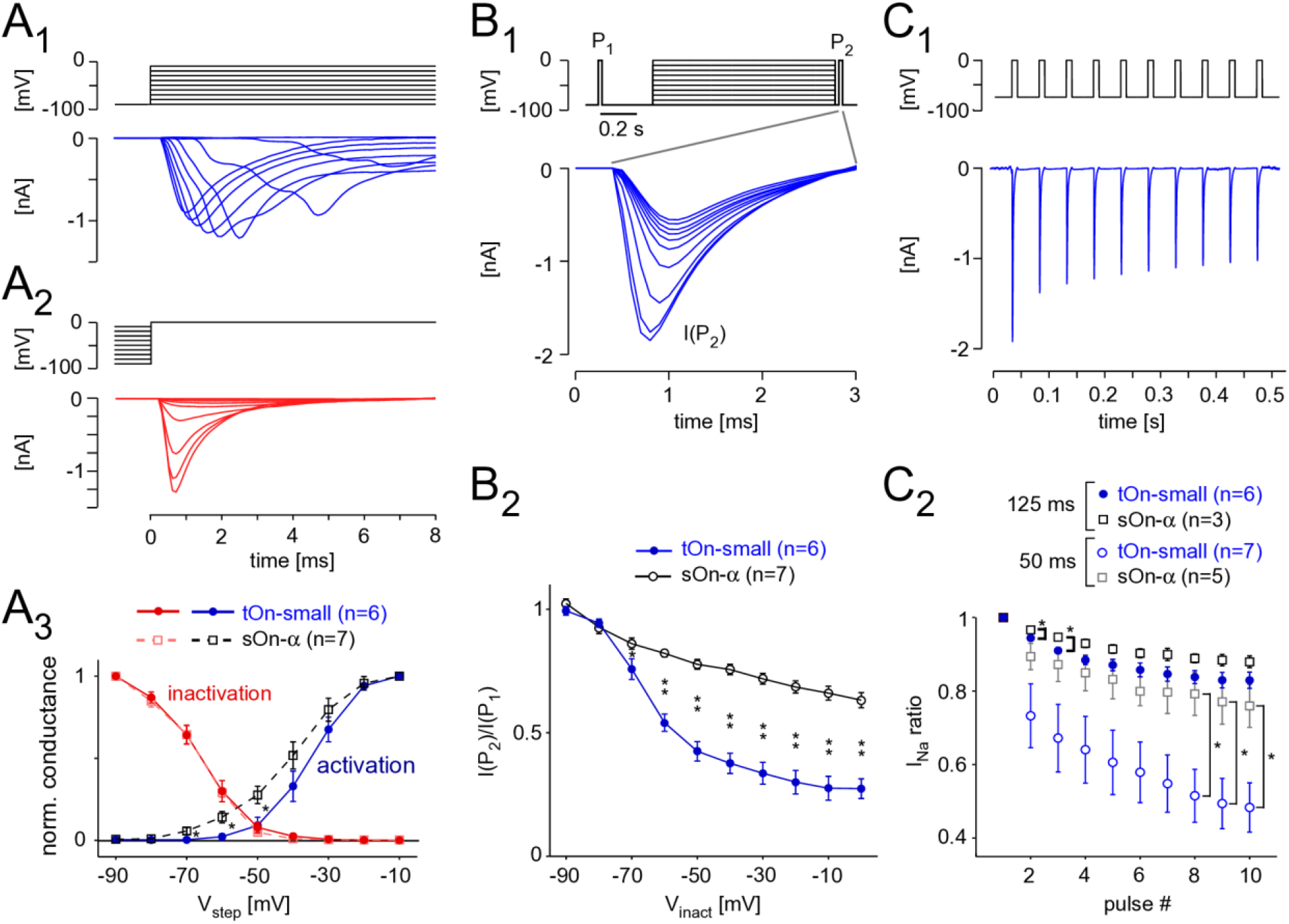
Voltage-gated Na^+^ channels in tOn-small cells desensitise strongly. **A**, Voltage-clamp recording of a tON-small cell to step protocols probing activation (A_1_) and inactivation (A_2_) of voltage-gated Na^+^ channels (VGSCs; A_1_, from -90 mV for 1 s to different voltages (−90 to -10 mV); A_2_, from different potentials (−90 to -10 mV) for 1 s to 0 mV; ΔV=10 mV for both protocols). A_3_, Normalized conductance as functions of step voltage (V_step_) for channel activation (squares) and inactivation (circles) in tOn-small (blue, n=6) and sOn-α cells (black, n=7). **B**, Probing inactivation of VGSCs with 3-ms test pulses (to 0 mV) delivered before (P_1_) and after (P_2_) the 1-s inactivating steps (B_1_, top; -90 to 0 mV, ΔV=10 mV, followed by an inactivating step for 20 ms to -90 mV). Na^+^ currents elicited by the 2^nd^ test pulse (P_2_) in a tOn-small cell (B_1_, bottom). B_2_, Ratio of Na^+^ current amplitudes elicited by the two test pulses (cf. B_1_) plotted against inactivating voltage. **C**, Probing “inactivation sensitivity” using sequences of 10-ms depolarizing voltage pulses (−70 to 0 mV). C_1_, Na^2+^ currents elicited in a tOn-small cell by a 20-Hz sequence. C_2_, Peak Na^+^ currents as a function of pulse number for 20 (Δt=50 ms) and 8 Hz (Δt=125 ms (blue circles and black squares for tOn-small and sOn-α cells, respectively). Peak Na^+^ currents were normalized to the amplitude evoked by the first pulse. Data from tOn-small (circles, n=7) and sOn-α cells (squares, n=5) for 20 Hz pulses, and tOn-small (circles, n=6) and sOn-α cells (squares, n=3) for 8 Hz pulses. Student’s t-test was used for determining statistical significance (*= p<0.05, **= p<0.01) between different conditions in A_3_, B_2_ and C_2_.

**Figure 6.**
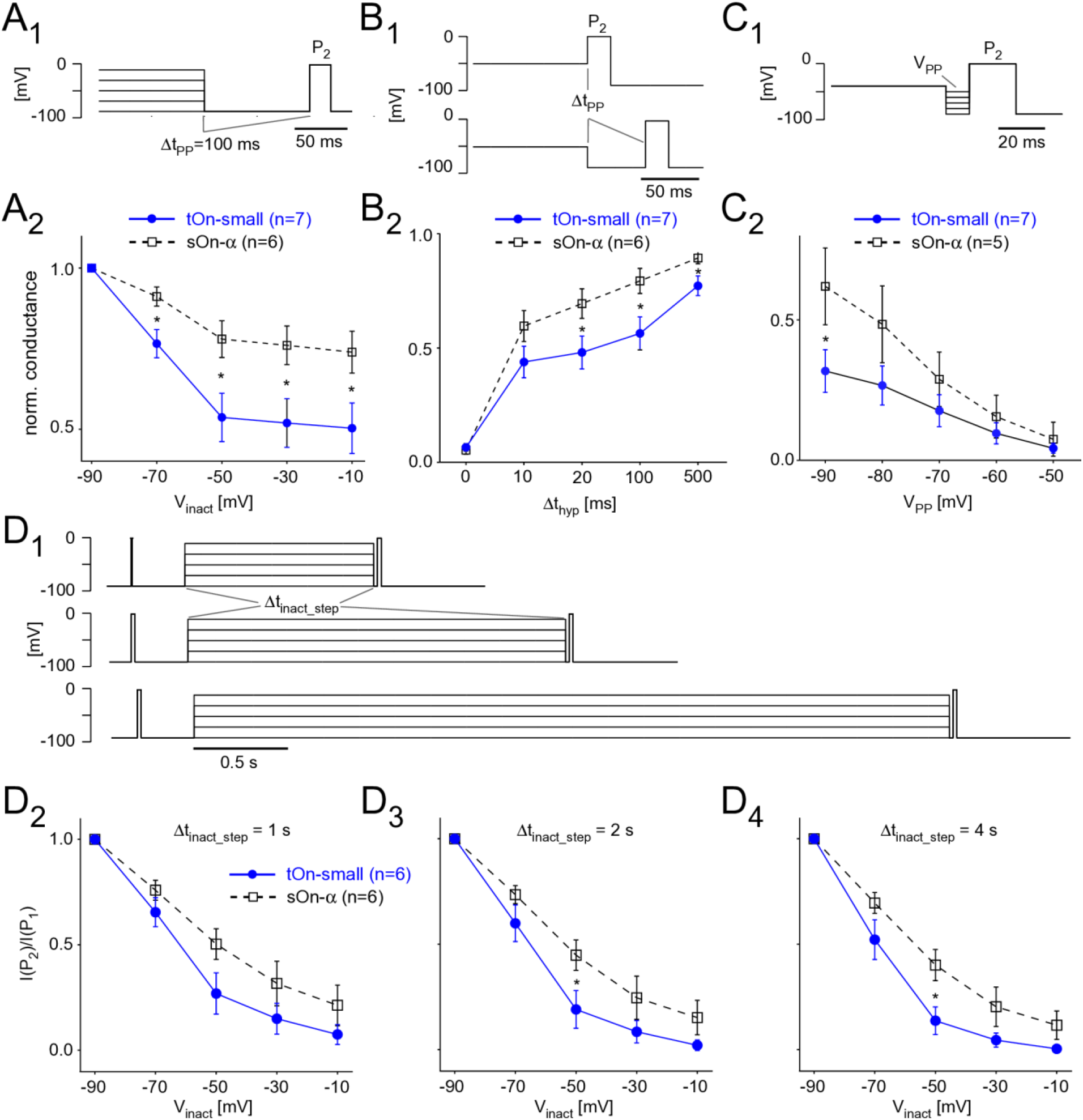
Steady-state slow inactivation of VGSCs shapes tOn-small RGC responses. **A**, Same protocol as Fig. 5B, but with a 100-ms hyperpolarizing pre-pulse (V_PP_ = -90 mV) inserted between inactivation step and test pulse P_2_; P_1_ not shown (A_1_). Normalized conductance as a function of inactivation step voltage for tOn-small (filled circles) and sOn-α RGCs (open squares; A_2_). **B**, Like in (A) but with pre-pulse duration varied (Δt_hyp_=0, 10, 20, 100, and 500 ms) for a single inactivation step voltage (V_inact_ = -50 mV). **C**, Like in (A) but with fixed Δt_hyp_= 10 ms and V_inact_= -40 mV, while varying pre-pulse voltage (V_PP_ = -90 to -50 mV). **D**, Same protocol as Fig. 5B but with the duration of the inactivation step (Δt_inact_step_) varied (D_1_). Test pulse current ratio (I(P_2_)/I(P_1_)) as a function of V_inact_for Δt_hyp_= 1 (D_2_), 2 (D_3_), and 4 s (D_4_). Student’s t-test was used for determining statistical significance (*= p<0.05) between different conditions in A-D.

### Voltage-gated Na^+^ channels in tOn-small RGCs need stronger hyperpolarization and longer times to recover from inactivation

It is known that VGSCs can undergo inactivation at different timescales and that their recovering from different states of inactivation may differ in its time- and voltage-dependence (e.g. (Martiszus et al., 2021; Tsai et al., 2011)). Hence, we next wanted to look at these aspects in the two cell types. To this end, we probed the cells with two test pulses (P_1_, P_2_), with P_2_ following a 1 s “inactivating” voltage-step and a short 20 ms hyperpolarizing pre-pulse (Fig. 5B_1_). While the previous protocol (Fig. 5A_2_) tested the combined effects of fast and slow inactivation, the insertion of a pre-pulse enables VGSCs to recover from fast inactivation and, hence, probing the effect of slow inactivation (Kim and Rieke, 2003; Silva, 2014). In the presence of this pre-pulse, the inactivating step reduced the test pulse response in both RGC types, but much more so in tOn-small cells (Fig. 5B_2_). This indicates that the VGSCs’ slow inactivation differed between the two cell types with respect to voltage-dependence, with the VGSCs in tOn-small displaying consistently stronger inactivation for step voltages ≥-70 mV.

To probe the time-dependence of recovery from inactivation, we first applied a pulse-train protocol with two different frequencies (Fig. 5C_1,2_), mimicking the activation by spikes (Kim and Rieke, 2003). We quantified VGSC inactivation by calculating the ratio between the Na^+^ current activated by the i^th^ pulse (*I*_*Na(i)*_) and that activated by the 1^st^ pulse of a train (*I*_*Na*(1))_. We found that *I*_*Na*_ ratio (*I*_*Na(i)*_/*I*_*Na*(1)_) in tOn-small cells dramatically dropped at 20 Hz already for the 2^nd^ pulse to ∼75% and for the 10^th^ pulse to less than 50%, whereas in sOn-α cells, *I*_*Na*_ ratio was only slightly reduced by consecutive pulses (Fig. 5C_2_). This indicates that VGSCs in tOn-small cells need substantially more time to recover from inactivation compared to those in sOn-α cells.

To study voltage- and time-dependence more closely, we went back to the two-pulse protocol. First, we varied the duration of the hyperpolarizing pre-pulse (Δ*t*_*PP*_) between inactivation step and second test pulse to determine the time-dependence of recovery from slow inactivation (Fig. 6A,B). With Δ*t*_*PP*_ = _0_, inactivation was virtually the same for both cell types (*cf*. Fig. 5A_2,3_), but became significantly different for Δ*t*_*PP*_ ≥ 20 *ms* before starting to approach similar values for Δ*t*_*PP*_=500 *ms* (Fig. 6B_2_), which supports a difference in time-dependence of recovery from slow inactivation. Next, we kept the duration of the pre-pulse constant but varied its voltage (V_*PP*_) to test the voltage-dependence (Fig. 6C_1_). We found tOn-small cells recovered only to approx. half the levels compared to sOn-α cells for all tested V_*PP*_ values (Fig. 6C_2_), arguing for an additional difference in voltage-dependence of recovery. Finally, we changed the duration of the inactivating step (Δ*t*_*inact*_ of 1, 2 or 4 s; Fig. 6D_1_) to make sure the difference between the cell types were not due to insufficient inactivation of the sOn-α cells; this seemed to have not been the case, as for all tested inactivation durations, tOn-small cells consistently showed stronger inactivation than sOn-α cells (Fig. 6D_2-4_).

Taken together, the differences observed in the spiking behaviour of tOn-small vs sOn-α cells likely arise from VGSCs in tOn-small cells (*i*) activate at a higher threshold, (*ii*) need stronger hyperpolarization and (*iii*) more time to recover from steady-state slow inactivation. These results support that idea that properties of the spike generator (i.e. of the VGSCs) importantly contribute to the temporal coding properties observed in tOn-small RGCs.

## DISCUSSION

Here we studied the underlying cell-intrinsic mechanisms that encode membrane voltage into spike patterns in two distinct types of RGC in the mouse retina. Recordings of light- and current-elicited voltage responses showed differences in the relationship of membrane voltage and elicited spikes between tOn-small and On-α cells: Unlike in sOn-α RGCs, spiking but not the voltage signal in tOn-small RGCs was very transient and high-contrast selective. Modelling and voltage-clamp recordings revealed that this response pattern of tOn-small cells is largely shaped by stronger desensitisation of the cell’s spike generator, i.e. its VGSCs. Our results support that notion that – in complement with upstream mechanisms and as the last step of the signal retinal processing chain – the intrinsic properties of an RGC can importantly shape the retina’s output to the brain.

### Multiple mechanisms can shape RGC response kinetics

RGC response properties can be shaped at different stage of retinal signal processing. First, RGCs can directly inherit diverse temporal response properties from their excitatory presynaptic partners, the BCs. It has been shown that transient BC types such as BC types 5t, 5o, 5i and XBC, mostly stratify in the central bulk of the IPL (Baden et al., 2013), therefore RGCs may become transient simply by having their dendrites tap into the respective IPL layers to collect transient excitatory input. Interestingly, the tOn-small cells likely correspond to the “6sn” cells classified in an EM dataset (Bae et al., 2018), which stratify slightly more towards the centre of the IPL than the sOn-α cells and thus can generate synapses with more transient BC types. This may contribute to more transient responses in tOn-small RGCs and more sustained responses in sOn-α cells. Second, inhibitory AC circuits can sharpen the time course of RGC responses either by shaping the signals at BC axon terminals or RGC dendrites (reviewed in (Diamond, 2017; Zhang and McCall, 2012)). Third, RGCs can employ intrinsic properties – i.e. kinetics of specific glutamate receptors and other postsynaptic channels, dendritic morphology and properties of the spike generator – to shape their spiking output. For example, in suppressed-by-contrast RGCs a depolarization block resulting from low VGSC conductance, short axonal initial segment (AIS) and selective expression of Na_v_ isoforms (see below) underlies the cells’ response (Wienbar and Schwartz, 2022). In complement, our study suggests that the functional properties of VGSCs expressed in tOn-small cells are the key mechanism that shapes these cells’ responses.

### Spike generator desensitisation and functional consequences

Our results suggest that desensitisation of the spike generator, likely based on slow inactivation of the VGSCs themselves, shapes spike coding in tOn-small cells in two ways: their responses are more transient and their sensitivity for low contrast stimuli is decreased compared with sOn-α cells. In previous studies, it was shown that slow inactivation of VGSCs participates in the adaptation of RGCs to temporal contrasts (Kim and Rieke, 2003; Weick and Demb, 2011). Another study investigating contrast adaptation in RGCs revealed two types of “plasticity” mechanisms: adaptation and sensitisation (Kastner and Baccus, 2011). Interestingly, the plasticity of the adapting type supports encoding of strong stimuli (i.e. high-contrast), whereas plasticity of the sensitising type promotes encoding of weak stimuli (i.e. low-contrast) – very similar to what we found in tOn-small (high contrast-selective) and sOn-α cells (also sensitive to low-contrast). Therefore, it is likely that desensitisation of the spike generator plays a functionally crucial role in at least three different aspects of RGC signalling: temporal kinetics (transient vs. sustained), contrast adaptation, and limiting the dynamic range for contrast encoding.

In our model, we simulated spike generator desensitization by changing the voltage dependence of the gating functions in a history-dependent manner (Methods). This can be thought as equivalent to slowing the recovery of the VGSCs from inactivation. Also, our voltage-clamp data points at the VGSCs to be involved in defining the response properties of tON-small cells: compared to sOn-α cells, the Na^+^ currents in tOn-small cells exhibited a higher threshold and recovered more slowly from inactivation. Therefore, that tOn-small and sOn-α cells likely differ in their VGSC complement.

Using RT-PCR, four VGSC α-subunit isoforms have been identified in rodent RGCs: Na_v_1.1, 1.2, 1.3 and 1.6 (Fjell et al., 1997). Among other things, these isoforms differ in their persistent current (reviewed in (Goldin, 1999)): For example, for more positive membrane potentials, the persistent current of Na_v_1.6 increases, whereas that of Na_v_1.1 decreases. A persistent current that increases with depolarization “pulls” the membrane potential towards the spiking threshold and, thus, may favour sustained spiking. Indeed, a recent study provided evidence that Na_v_1.6 is dominantly expressed in mouse sustained OFF alpha RGCs, whereas suppressed-by-contrast RGCs likely express a different isoform (Wienbar and Schwartz, 2022). Thus, the expression ratio of VGSC isoforms affects the time course of an RGC’s spiking response. In addition, VGSCs are strongly modulated by accessory β-subunits (reviewed in (Goldin, 1999)), which can, for example, slow or accelerate inactivation, and shift voltage-dependence. Therefore, not only VGSC properties but also differential β-subunit expression may also contribute to the differences we observed between tOn-small and sOn-α cells.

Additionally, it was shown that a small subset of RGCs additionally expresses an unusual TTX-insensitive VGSC α-subunit isoform in the soma and the proximal dendrites: Na_v_1.8 (O’Brien et al., 2008). These cells had large somata and were neurofilament-positive, suggesting that alpha (but not tOn-small) cells express Na_v_1.8. In contrast to the above VGSC isoforms, Na_v_1.8 mediates currents that exhibit very little subthreshold inactivation and recover much faster from inactivation (Cummins and Waxman, 1997; Renganathan et al., 2000). It is possible that channels with such properties contribute to the high contrast sensitivity – for instance, by enabling dendritic spiking (Velte and Masland, 1999) – and the sustained spiking response of sOn-α cells. Nevertheless, we did not observe differences in inactivation between tOn-small and sOn-α cells which argues against a prominent role of Na_v_1.8 in sOn-α cells – at least under our experimental conditions.

Finally, VGSCs are precisely arranged in type-specific subdomains along the AIS (Van Wart et al., 2007). These VGSC “bands” appear to vary in length and location (relative to the soma) between RGC types (Fried et al., 2009; Wienbar and Schwartz, 2022). It was suggested that this organization shapes the response properties of the RGC spike generator, including the activation threshold (Fried et al., 2009). If the VGSC bands along the AIS of tOn-small and sON-α cells differ remains to be investigated.

### Diverse retinal computations are performed by intrinsic properties of RGCs

A great deal of neural computations taking place in the retina ultimately segregates features of the visual stimulus into diverse parallel channels to the brain (reviewed in (Baden et al., 2020; Gollisch and Meister, 2010; Kerschensteiner, 2022; Wassle, 2004)). Complex interactions involving different sets of interneurons in the two plexiform layers of the retina contribute to this parallel feature extraction (reviewed in (Diamond, 2017; Franke and Baden, 2017)). Our study highlights that a mechanism at the last step of retinal processing – intrinsic properties of the RGCs – can strongly participate in forming their responses. This is in line with several previous studies showing that spike generation in RGCs is rather diverse in its location and dynamics (see also above): For example, it was reported that Na^+^-based spikes can originate from both the dendrites and the soma region of RGCs (Oesch et al., 2005; Velte and Masland, 1999) and dendritic spikes play an important role in sharpening the directional tuning of DS RGCs (Oesch and Taylor, 2010; Schachter et al., 2010; Velte and Masland, 1999). Moreover, RGCs display very different levels of spike frequency adaptation in response to current injection (O’Brien et al., 2002). These studies, together with the present one, support the notion that RGCs are not the passive integrators of upstream signals – as they are still depicted in textbooks – but play an active role in retinal processing.

Still, the question arises what the advantage of feature extraction at the very last level of retinal processing may be. Considering the “feature” that is extracted by the tOn-small RGC – selective high-contrast signals – one may speculate that this RGC type exclusively relays very strong and reliable events to higher visual areas. Extracting this feature late in the retinal network may bear the advantage that signal-to-noise is at its maximum at the RGC level, providing a suitable high-quality signal to cut off low-contrast responses. Additionally, a feature extraction mechanism intrinsic to RGCs would be largely independent from the highly dynamic inner retinal circuits and could provide “stability” to the output signal. In light of evidence suggesting that the retinal code changes over the full range of illumination conditions (Tikidji-Hamburyan et al., 2014), a spike generator with properties like the one in tOn-small RGCs may provide a “stable” element for encoding high-contrast signals across different visual environments.

## MATERIALS AND METHODS

### Animals and tissue preparation

The Tg(Igfbp5-EGFP)JE168Gsat transgenic mice were obtained from the Mutant Mouse Regional Resource Centre (MMRRC; University of California, Davis, CA, USA). In this transgenic mouse line, the EGFP reporter gene, followed by a polyadenylation sequence, was inserted into the BAC clone RP24-159O10 at the initiating ATG codon of the first coding exon of the Igfbp5 gene so that EGFP expression is driven by the regulatory sequences of the BAC gene (Gong et al., 2003). The resulting modified BAC (BX1812) was used to generate this transgenic mouse line (The Gene Expression Nervous System Atlas [GENSAT] Project, The Rockefeller University, New York, NY), to which we refer in the following as “*Igfbp5*” line.

All procedures were performed in accordance with the law on animal protection issued by the German Federal Government (Tierschutzgesetz) and approved by the institutional animal welfare committee of the University of Tübingen or the MPI for Brain Research, Frankfurt/M. Mice of both genders (4-8 weeks of age) were housed under a standard 12 hr day/night rhythm. For electrical recordings, mice were dark-adapted for ≥1 h before the experiment. The animals were then anesthetized with isoflurane (Baxter) and killed by cervical dislocation.

For immunostainings, the mouse eyes were dissected in cold 0.1 M phosphate buffer (PB), pH 7.4 and the posterior eyecups were immersion fixed in 4% paraformaldehyde (PFA) in PB for 15–30 min at room temperature. Following fixation, retinas were dissected from the eyecup, cryo-protected in graded sucrose solutions (10, 20, 30% w/v), and stored at –20°C in 30% sucrose until use. Retinal pieces were sectioned vertically at 16-20 µm using a cryostat.

For electrical recordings, the mouse eyes were enucleated and hemisected in carboxygenated (95% O_2_, 5% CO_2_) artificial cerebral spinal fluid (ACSF) solution, which contained (in mM): 125 NaCl, 2.5 KCl, 2 CaCl_2_, 1 MgCl_2_, 1.25 NaH_2_PO_4_, 26 NaHCO_3_, 0.5 L-glutamine, and 20 glucose; maintained at pH 7.4. After removal of the vitreous body, each retina was flat-mounted onto an Anodisc (#13, 0.2 um pore size, GE Healthcare, Maidstone, UK) with the ganglion cell layer facing up. The Anodisc with the tissue was then transferred to the recording chamber of a two-photon (2P) microscope or a Zeiss Axioscope (see below), where it was continuously perfused with carboxygenated ACSF. When using the 2P microscope, ACSF contained 0.5–1 µM Sulforhodamine 101 (SR101, Invitrogen Steinheim, Germany) to reveal blood vessels and any damaged cells in the red fluorescence channel (see below). All procedures were carried out under very dim red (>650 nm) light.

### Antibodies and immunohistochemistry

Cholinergic amacrine cells were labelled with goat anti-choline acetyltransferase (ChAT, 1:200; Chemicon), GABAergic amacrine cells with rabbit anti-GABA (1:2000; Sigma). A rat anti-glycine antibody labelled all glycinergic amacrine cells and ON cone bipolar cells (1:1000, kindly provided by David Pow, Brisbane, Australia). Rabbit anti-GFP was used to increase the EGFP signal (1:2000, Molecular Probes). The neurofilament marker SMI32 (mouse monoclonal, 1:1000; Sternberger Monoclonals) and a goat anti-osteopontin antibody (OPN, 1:1000; R&D Systems) were used to label alpha RGCs. Immunocytochemical labelling was performed using the indirect fluorescence method. Sections were incubated overnight with primary antibodies in 3% normal donkey serum (NDS), 0.5% Triton X-100, and 0.02% sodium azide in PB. After washing in PB, secondary antibodies were applied for 1 h. These were conjugated either to Cy3 (Dianova), or Alexa Fluor488 (Invitrogen).

Confocal images were taken by using a Zeiss LSM 5 Pascal confocal microscope equipped with an argon and a HeNe laser. Images were taken with a Plan-Neofluar 40x/1.3 objective. Figures represent projections calculated from stacks of images with the LSM software or ImageJ (W.S. Rasband, http://imagej.nih.gov/ij/). Brightness and contrast of the final images were adjusted using Adobe Photoshop.

### Single cell injection with DiI

For dye injections of bipolar cells, enucleated eyes were transferred to oxygenated Ames medium (Sigma-Aldrich, Taufkirchen, Germany) and opened by an encircling cut. The retinas were dissected and embedded in 2% low melting agar (2-hydroxymethyl agarose, Sigma Aldrich), mounted on a vibratome (DSK Microslicer, DTK-1000, Ted Pella, Inc), and cut into 150 µm sections. After another 10 min in Ames Medium, sections were fixed in 4% PFA in PB at 4°C for 15 min. For injections with the fluorescent lipophilic tracer DiI (Molecular Probes) sharp microelectrodes were pulled from borosilicate glass tubing (Hilgenberg, Malsfeld, Germany) and filled with 0.5% DiI solution in 100% ethanol. The dye was injected into EGFP-labelled bipolar cells with 1nA positive current for 3 min. For DiI injections of amacrine cells and ganglion cells, retinal whole mounts were fixed in 4% PFA in PB for 15-30 min. After fixation, the tissue was kept in PB at 4°C overnight for the efficient diffusion of dye into the fine dendrites and axons. The filled cells were documented by using the confocal microscope.

### Two photon microscopy

We used a MOM-type two-photon microscope (designed by W. Denk, MPImF, Heidelberg; purchased from Science Products/Sutter Instruments, Novato, USA). Both design and procedures were described previously (Euler et al., 2009). The system was equipped with a mode-locked Ti:Sapphire laser (MaiTai-HP DeepSee, Newport Spectra-Physics, Germany) tuned to 927 nm, two detection channels for fluorescence imaging (red, HQ 622 BP36; green, D 535 BP 50, or 520 BP 39; AHF, Tübingen, Germany) and a 20x objective (XLUMPlanFL, 0.95 NA, Olympus, Hamburg, Germany). The red fluorescence channel was used to visualize the retinal structure using SR101 staining (see above), the green channel to target EGFP-labelled *Igfbp5* RGCs.

### Electrophysiology

In the *Igfbp5* line, two types of RGCs were targeted for intracellular recordings: Non-fluorescent sOn-α cells, identified by their very large polygonal somata, and EGFP-labelled tOn-small RGCs, which possess medium-sized somata (∼15 μm in diameter). The identity of the recorded RGC type was confirmed after the recording based on the dendritic morphology. To avoid confounds related to morphological variability across the retina (Bleckert et al., 2014), the cells were collected (and recorded) from the ventral retina (∼0.8 mm from the optic disc).

For recording of light responses (current-clamp), electrodes (with resistances of 5–10 MΩ) contained (in mM): 120 K-gluconate, 5 NaCl, 10 KCl, 1 MgCl_2_, 1 EGTA, 10 HEPES, 2 Mg-ATP, and 0.5 Tris-GTP, adjusted to pH 7.2 using 1M KOH. In addition, 4% Neurobiotin (Molecular Probes, Eugene, USA) and 0.2 mM SR101 were added to reveal the cell’s dendritic morphology. Membrane voltage was corrected for a liquid junction potential of ∼14 mV. Data were acquired using an Axoclamp-900A amplifier (Molecular Devices GmbH, San Jose, USA) and digitized at 10 kHz. Experiments were carried out at 37°C. After the electrical recordings, the tissue was fixed and cells were visualized by overnight incubation in 1:1000 Streptavidin-Alexa Fluor 594 (Invitrogen, Darmstadt, Germany). The RGC’s morphology (as image stack) was documented and Z-projection images were made using ImageJ.

For voltage-clamp recordings, CdCl_2_ (100 µM) was added to the ACSF to block currents through voltage-gated Ca^2+^ channels. Electrodes (with resistances of 5-8 MΩ) contained (in mM): 120 Cs-gluconate, 1 CaCl_2_, 1 MgCl_2_, 10 Na-HEPES, 11 EGTA, 10 TEA-Cl, Sulforhodamine B (0.005%), adjusted to pH 7.2 with CsOH. Liquid junction potentials of 15 mV were corrected before the measurement with the pipette offset function of the amplifier. Series resistance ranged from 5 to 21 MΩ (13.7 ± 8.3 MΩ, n=25) and cell capacitance were not compensated. Seal resistances >2 GΩ were routinely obtained. Cells were voltage-clamped at -90 mV and the different step protocols applied (see Figs. 5,6). All experiments were carried out at room temperature (20-22°C). Data were acquired using an Axopatch-200B amplifier (Molecular Devices, San Jose, USA), digitized at 10 kHz using the pClamp software (Molecular Devices), and Bessel-filtered at 2 kHz.

### Light stimulation

For light stimulation, we used a small reflective liquid-crystal-on-Silicon (LCoS) display (i-glasses; EST), coupled in the microscope’s optical path. The display was alternately illuminated by two bandpass-filtered (blue, 400 BP 10; green, 578 BP 10; AHF) LEDs, projecting spatio-temporally structured stimuli through the objective lens onto the retina (Euler et al., 2009). Stimulus intensity was measured using a calibrated photometer (Model 842-PE, 200-1100 nm, Newport) set to the respective centre wavelength of the LED filters. Cone photo-isomerization rates were calculated as described previously (Chang et al., 2013); all stimuli featured equal photo-isomerization rates for M- and S-opsin. Five stimulus protocols were used:

(a) a 200 µm-diameter bright spot flashed for 1 s,

(b) a 300 × 1000 µm bright bar moving at 500 µm/s in 8 directions,

(c) a 1 Hz sinusoidally-modulated bright spot of varying diameter at 57% mean contrast,

(d) a 1 Hz sinusoidally-modulated spot of varying contrasts, with the optimal spot diameter determined by stimulus (c),

(e) a flickering (white noise) spot, with the optimal spot diameter determined by stimulus (c); spot intensity was randomly chosen from a binary distribution for each frame (80 Hz refresh rate) and the mean intensity equal to background intensity.

For sinusoidally modulated stimuli (c, d), contrast was defined as the ratio of s.d. and mean intensity. The retina was always illuminated with a constant background in the (low) photopic range. In case of the protocols a and b, background illumination generated photo-isomerization rates (in 10^4^·P*s^-1^ per photoreceptor) of 1.2, 0.9 and 2.2, and in case of protocols c to e, 2.3, 2.2 and 4.8 for M-opsin, S-opsin and rhodopsin, respectively.

## Data Analysis

All electrophysiological data were analysed off-line using custom MATLAB scripts (Mathworks, Ismaning, Germany). To analyse *tOn-small* RGC responses to moving bars (protocol b), we defined the stimulus direction that generated most spikes (calculating the vector sum of the total spiking responses in 8 directions) as “preferred direction”, and calculated a direction selective index (*DSi*) as follows:

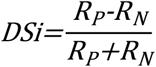

with preferred direction-response *R*_*P*_ and null (=opposite) direction response *R*_*N*_.

For the white noise stimulus (protocol e), we used the linear-nonlinear (LN) cascade model described earlier (Baccus and Meister, 2002; Chichilnisky, 2001; Field et al., 2010; Kim and Rieke, 2001; Wang et al., 2011) to interpret RGC responses. This model consists of a linear filter that determines the cell’s temporal, chromatic and spatial sensitivities, as well as a “static” nonlinearity that converts the filtered stimulus into a firing rate. In the time domain, the linear filter is proportional to the spike-triggered average stimulus (STA, the average stimulus preceding each spike) (Chichilnisky, 2001). Therefore, for LN models with identical linear filter but different nonlinearities, spike-triggered average stimuli are identical up to a scale factor. We estimated the “static” nonlinearity from the nonlinear relationship between linear filtered responses (“generator signal”) and real responses.

Due to the limited refresh rate of white noise stimulus (80 Hz), we defined number of spikes in a time bin of Δ*t* = 12.5 ms (1/80 Hz) as the cell’s response, and STA was computed as the cross-correlation between response and stimulus. For graded signal, the cell’s response was defined as relative average voltage within each time bin (by subtracting the minimum voltage of the recorded trace).

We used the Difference-of-Gaussians (DOG) receptive field model in two dimensions to describe the spatial structure of the recorded RGCs. We assumed that the receptive field (RF) of each cell could be approximated by the weighted difference of two concentric Gaussian, one for the RF centre and one for the RF surround. A nonlinear function was included into the model to convert this weighted sum into the firing rate (Yin et al., 2009). An RGC’s response *R* to a spot stimulus centred in the RF and with diameter *D* and contrast *c* was calculated as follows:

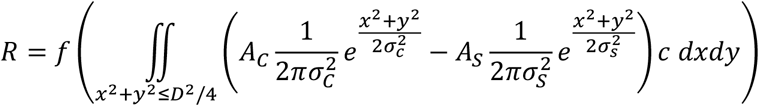

with the nonlinear function *f*, amplitude (*A*_*C*_) and spatial extent (*σ*_*C*_) of the RF centre, as well as amplitude (*A*_*C*_) and spatial extent (*σ*_*S*_) of the RF surround.

The nonlinear function *f* was determined using the same RGC’s responses to spots with fixed diameter but varying contrasts, fitted by the Naka-Rushton Equation (Naka and Rushton, 1966):

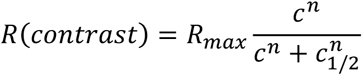

The other four parameters (*A*_*C*_, *A*_*S*_, *σ*_*C*_, *σ*_*S*_) were determined by fitting the model to area summation data using a numerical search.

### Computational modelling

To explain our experimental data, we constructed an one-compartment Hodgkin-Huxley model,

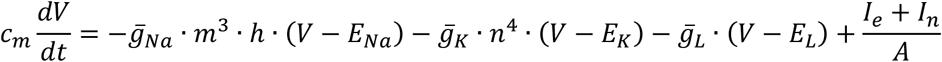

with the specific membrane capacitance *c*_*m*_ = 10 nF/mm^2^, the voltage across the cell membrane (*V*) in mV, the maximal conductance for transient Na^+^ current 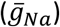, delayed-rectifier *K* ^+^ current and a leakage current 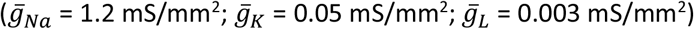, the reversal potentials for the three currents (*E*_*Na*_ = 50 mV; *E*_*K*_= -76 mV; *E*_*L*_= -70 mV), the area of the cell membrane (*A* = 0.0013 mm^2^), the injected current (*I*_*e*_), and a noise component (*I*_*n*_) with a 1/*f* power spectrum (pink noise). In Fig. 4C, *I*_*e*_ is the estimated response to a sequence of binary white noise, using the linear filter and static nonlinearity determined from graded voltage responses (Fig. 4A); in Fig. 4D, *I*_*e*_ are current steps with varying amplitudes. As gating functions for *m, h*, and _*n*_ we used:

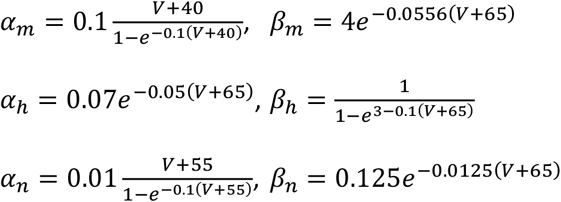

After each spike, the voltage dependence of the gating functions for *m* and *h* was increased by 1.55 mV for the sensitising model (i.e. tOn-small RGCs) and by 0.01 mV for the non-desensitising case (i.e. sON-α RGCs). The recovery from this shift in voltage dependence followed an exponential function with a time constant of 5 s.

## Acknowledgements

This study was funded by Deutsche Forschungsgemeinschaft (DFG) (HA 5277/3-1, EU 42/4-1 and FOR 701). We thank Tom Baden, Philipp Berens, and Katrin Franke for helpful discussions and Gordon Eske for technical assistance.

## EXTENDED DATA

**Figure 1-1.**
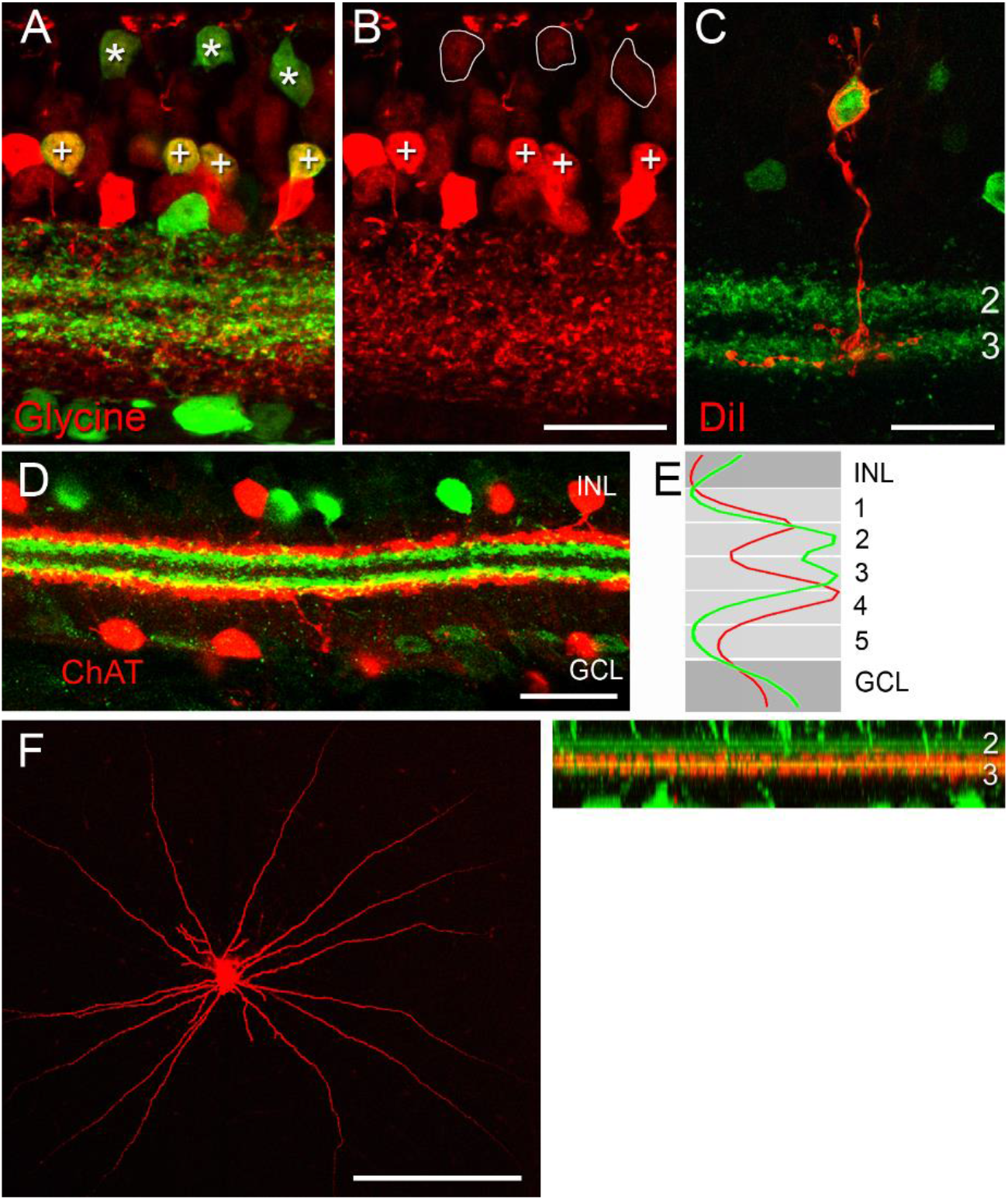
EGFP-labelled bipolar and amacrine cells in the *Igfbp5* transgenic mouse line (related to Fig. 1) **A**,**B** The INL contained many EGFP-labelled cells, including some GABA-positive ACs (data not shown) and many glycine-positive cells. The latter were comprised of ACs and On cone BCs. Vertical section of *Igfbp5* retina double-labelled for EGFP (green) and glycine (red). EGFP-labelled ACs in the second-inner row of the INL (+) and ON BCs (*) contain glycine. EGFP-labelled processes extend along sublaminae 2 and 3 of the inner plexiform layer (IPL). **C**, DiI-injections (n=12 cells) revealed that the EGFP-labelled BCs morphologically corresponded to On cone BC type 5. Example of an individual DiI-injected EGFP-expressing BC with typical type 5 morphology, with axon terminals stratifying in sublamina 3. **D**, Vertical section double-labelled for EGFP (green) and ChAT (red). **E**, Fluorescence intensity profile along z-axis of a confocal stack of whole mount retina double-labelled for EGFP (green) and ChAT (red). **F**, Analogous dye injections showed that the EGFP ACs included various monostratified, medium- and wide-field cells with On or Off stratification (data not shown). Example of *Igfbp5*-positive amacrine cell, following DiI filling in a retinal wholemount (left) with dendrites (red) stratifying in sublamina 3 (right). Scale bars: **B-D**, 20 µm; **F**, 200 µm.

**Figure 1-2.**
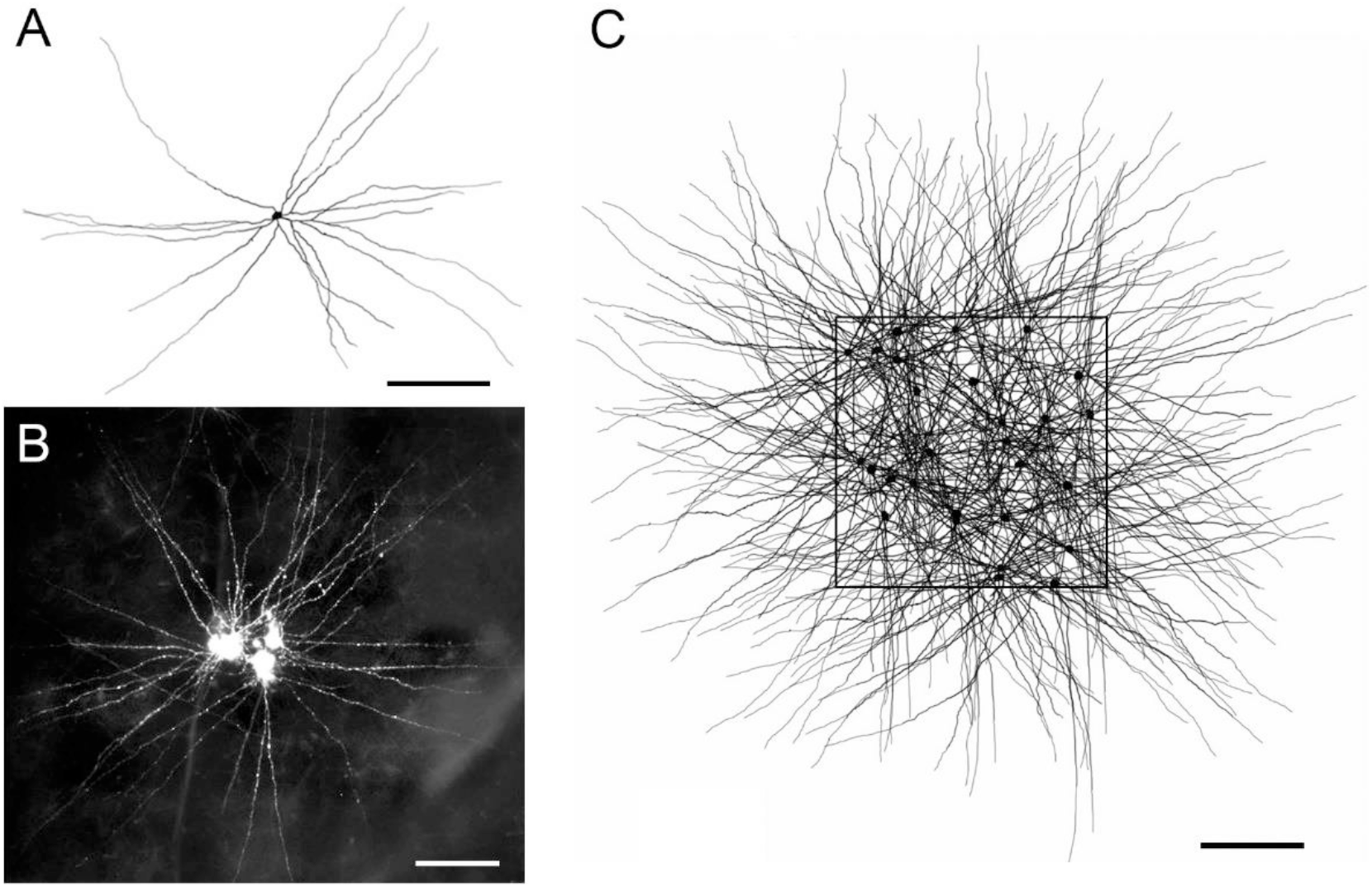
The mosaic of *Igfbp5*-positive On amacrine cells (related to Fig. 1) **A**, Reconstruction of a single, representative I*gfbp5*-positive amacrine cell, following DiI filling in a retinal wholemount. **B**, Mosaic of 5 neighbouring *Igfbp5*-positive amacrine cells filled with DiI. **C**, Illustration of the *Igfbp5* plexus serving the retinal patch delimited by the square. The patch contained 27 *Igfbp5*-positive amacrine cells (black square), to which 5 different representative morphological reconstructions were assigned randomly. Note that although very dense, the plexus is not complete, as dendrites from cell bodies located outside the depicted patch would cross this retinal area. Scale bars: **A-C**, 200 µm

**Figure 2-1.**
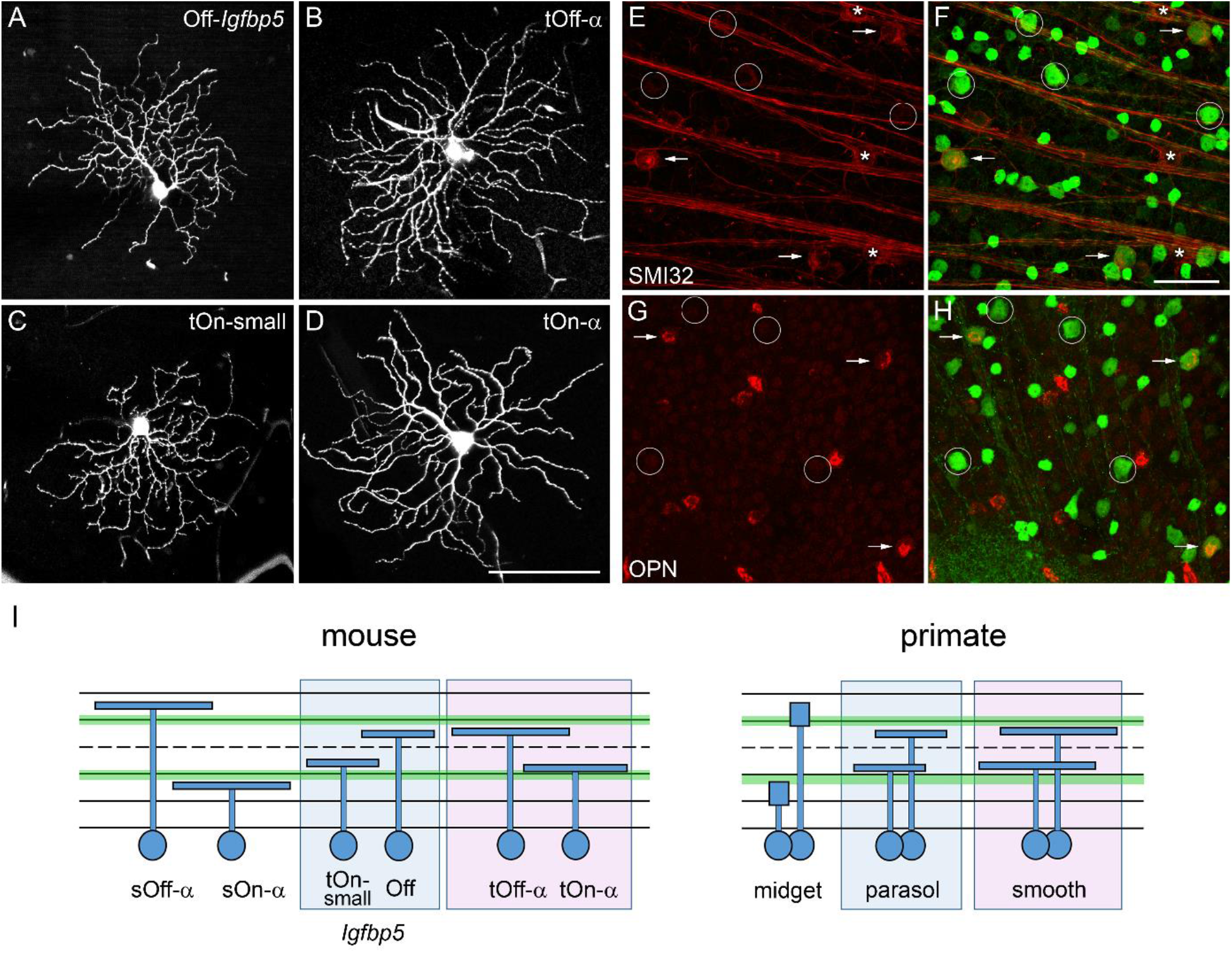
Retinal ganglion cells in the *Igfbp5* retina (related to Fig. 2) **A-C**, Examples of EGFP-positive RGCs: Off RGC (A), transient Off-alpha (tOff-α) RGC (B), and transient On small (tOn-small) RGC (C). **D**, For comparison, example of a (*Igfbp5*-negative) sustained On alpha (sOn-α) RGC. The cells were all collected (and filled with Neurobiotin) from the ventral retina (∼0.8 mm from the optic disc). **E-H**, On *Igfbp5*-positive RGCs are not immunoreactive for SMI32 and OPN. *Igfbp5* retina double-labelled with anti-neurofilament marker SMI32 (red) [1–3] and anti-GFP (green; E,F). sOn-α (asterisks) and tOff-α RGCs (white arrows) were SMI32-positive, whereas tOn-small RGCs (circles) were SMI32-negative. G,H, *Igfbp5* retina double labelled with anti-osteopontin (OPN, red) and anti-GFP (green; G,H). tOff-α RGCs are OPN-positive (white arrows), whereas tOn-small RGCs are OPN-negative (circles). **I**, Possible RGC homologues in mouse and primate retina: Dendritic stratification depth of mouse RGCs within the IPL (left): sOff-α and sOn-α RGCs, Off and tOn-small *Igfbp5*-positive RGCs, tOff-α and tOn-α RGCs. For comparison, dendritic stratification depth of primate RGCs (right): Off and On midget RGCs, Off and On parasol RGCs, and Off and On smooth RGCs (adapted from [4]). Scale bars: A-D, 100 µm; E-H, 50 µm.

**Figure 3-1.**
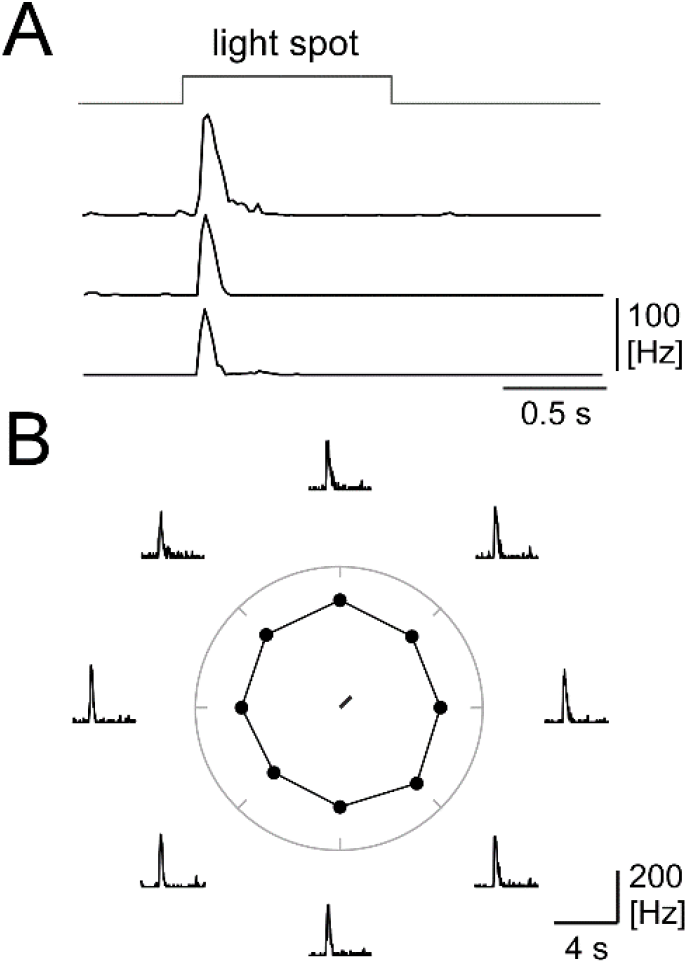
tOn-small ganglion cells are transient but not direction-selective (related to Fig. 3). **A**, Spike responses of three tOn-small RGCs to 1-s step light flash (diameter = 200 µm; average from 10 trials each, 20-ms time bins). **B**, Spike responses of an exemplary tOn-small cell to bar stimuli moving in 8 directions; responses shown as peri-stimulus time histogram (averages from 10 trials per direction, 20-ms time bins). Centre: Polar plot showing directional tuning curve and vector sum (calculated from the mean spike rate during stimulation for 8 directions) for an exemplary tOn-small RGC.

